# A highly scalable approach to topic modelling in single-cell data by approximate pseudobulk projection

**DOI:** 10.1101/2024.02.21.581497

**Authors:** Sishir Subedi, Tomokazu S Sumida, Yongjin P Park

## Abstract

Probabilistic topic modelling has become essential in many types of single-cell data analysis. Based on probabilistic topic assignments in each cell, we identify the latent representation of cellular states, and topic-specific gene frequency vectors provide interpretable bases to be compared with known cell-type-specific marker genes. However, fitting a topic model on a large number of cells would require heavy computational resources–specialized computing units, computing time and memory. Here, we present a scalable approximation method customized for single-cell RNA-seq data analysis, termed ASAP, short for Annotating Single-cell data by Approximate Pseudobulk estimation. Our approach is more accurate than existing methods but requires orders of magnitude less computing time, leaving much lower memory consumption. We also show that our approach is widely applicable for atlas-scale data analysis; our method seamlessly integrates single-cell and bulk data in joint analysis, not requiring additional preprocessing or feature selection steps.

## Introduction

### Background

High-throughput single-cell sequencing has gained popularity and has been successfully applied to many recent studies to understand cellular contexts. Not only the number of studies that involve single-cell sequencing but also the number of cells that a single study can profile has dramatically increased from hundreds to millions. Such technological advances have accompanied computational methods that can provide necessary operations in almost every step of data analysis–from data procurement to final statistical inference (Kharchenko 2021). However, typical computing infrastructures are often not well-equipped with powerful computing units and high-capacity memory despite the fact that many advancements in computational methods assume such large resources are readily accessible (Heumos et al. 2023; Zhang et al. 2024). Training a sophisticated model on millions of single-cell data vectors, using stochastic gradient descent algorithm, not surprisingly demands much more resources than fitting a model on data with thousands of cells and a thousand features.

After the quantification of molecular features within each cell, a conventional single-cell analysis (Stuart and Satija 2019; Stoeckius et al. 2017; Hao et al. 2021) generally turns to dimensionality reduction, such as principal component analysis (PCA) (Hotelling 1933; Jolliffe 1986), nearest neighbour graphs were constructed by matching cells based on the reduced data, and cell clusters are often resolved by a graph-based clustering method, such as Louvain (Blondel et al. 2008) or Leiden algorithm (Waltman and Eck 2013; Traag et al. 2018). The complexity of an exact PCA algorithm (singular value decomposition) linearly scales with the number of genes/features and quadratically with the number of cells (Tsuyuzaki et al. 2020), designing a scalable latent data modelling is crucially important (Sun et al. 2019). Should a large number of cells be analyzed, finding hidden gene regulatory programs/factors/topics is a computationally intensive task. It is customary to conduct an unvetted feature selection for highly variable genes to reduce memory footprint. By doing so, we critically assume that biologically relevant signals are most pronounced in data and further takes that the rank of gene expression variance of the selected genes is somewhat stable across different data sets.

### Related work

How can we uncover latent gene programs without compromising the intrinsic dimensionality? Several approaches have been suggested to overcome the scalability issues of the embedding or dimensionality reduction step before or after building neighbour graphs, and the ultimate outcome is to resolve clusters of cells using a graph-based clustering method. Most existing methods address the question of how we select fewer representative cells to ease the computational burden of downstream analysis, not sacrificing the accuracy of the final clustering results (Hie et al. 2019b; Abdelaal et al. 2020; Wei et al. 2022; Dhapola et al. 2022). A geometric sketching method was first coined to randomly sample cells while uniformly covering underlying and unknown metric space (Hie et al. 2019b). A similar idea was explored in natural language processing but more focused on defining the convex hull (boundary) of word occurrence topics over document corpus (Mimno and Lee 2014). More recent scalable approaches developed by the genomics community also build on the same premise that we can approximately capture the overall clustering patterns across millions of cells by choosing representative (anchor) cells wisely (Abdelaal et al. 2020; Wei et al. 2022; Dhapola et al. 2022). After building nearest neighbour graphs of the anchor cells, the rest of the algorithmic steps in all these approaches largely focus on adding new cells (vertices) to the original nearest neighbour graphs of the initial anchor cells and performing label propagation of cell type annotations.

### Our approach

Instead, here, we focus on developing a scalable approach for probabilistic topic modelling. We deal with high dimensionality and large sample size by mapping high-dimensional expression/activity vectors onto lower-dimensional topic space via random projection. Probabilistic topic modelling decomposes a large count data matrix into an interpretable dictionary matrix (topic-specific activities across tens of thousands of genes/features) and a topic proportion matrix (each cell’s attribution to the topics) (Blei et al. 2003; Dieng et al. 2020). Large-scale single-cell analysis can be made quite rapidly by considering the data generation process from a rather different perspective. Instead of designing a new type of complex model while leaving its statistical inference up to machine learning libraries, we propose a new framework in which a fundamental topic model can be estimated in an accurate, robust, and scalable way.

### Why topic modelling?

Fitting a probabilistic topic model has long been considered a principled and intuitive approach to uncovering patterns hidden underneath count data derived from high-throughput sequencing. A grade of membership model (or admixture) was first coined in genetics while trying to identify population structures manifested in genetics variant counts across the genome (Pritchard et al. 2000; Novembre 2016). In the same way, multi-tissue, multi-individual gene expression patterns were analyzed by the same type of model (Dey et al. 2017). More recently, Poisson matrix factorization (PMF) was shown to be an equivalent problem, and a more efficient method based on alternating regression estimation was suggested (Carbonetto et al. 2021, 2023). In single-cell genomics, especially for sparse DNA accessibility data, embedding methods based on topic modelling (or latent semantic indexing), such as ArchR (Granja et al. 2021) and cisTopic (Bravo González-Blas et al. 2019), were ranked in the top lists of a recent benchmark study (Chen et al. 2019). More recently, a topic model approach based on a deep variational autoencoder model, called embedded topic model, or ETM (Dieng et al. 2020), has been successfully used in single-cell RNA-seq modelling (Zhao et al. 2021b; Subedi and Park 2023; Zhang et al. 2023).

### Our work

We present a new scalable and versatile method that can quickly recover cellular topics with modest computing resources. We term our method ASAP, short for annotating a single-cell data matrix by approximate Psueudobulk estimation. Several existing works and observations inspired us. First, repeatedly applying random projection operation onto high-dimensional feature vectors may result in quick and moderately accurate cell clustering patterns (Wan et al. 2020). Second, pseudobulk data derived from aggregating within cell types and states often behave similarly to typical RNA-seq bulk data. Third, a grade of membership, or equivalent non-negative matrix factorization (NMF), method is powerful enough to dissect cellular topics from bulk sequencing profiles (Dey et al. 2017) and single-cell sequencing data (Carbonetto et al. 2021).

## Results

### Overview of ASAP

Briefly, ASAP will handle a matrix factorization problem of massive single-cell data in the following three steps: (1) Randomly project cells onto low-dimensional space (Fig 1A) to sort them through a binary classification tree (Fig 1B), depending on the sign of the random projection (RP) values, and construct a pseudobulk (PB) data matrix by aggregating cells landed in the same termini of the binary sorting tree. As suggested by the previous work (Wan et al. 2020) and demonstrated in the UMAP (Becht et al. 2018) (uniform manifold approximation and projection), simple RP operations can help cells with similar expression variation group together (Fig 1B). It is also encouraging to find that the resulting PB data already show that cells of similar cell types are naturally enriched within the same PB sample (the structure plot of Fig. B, cell type proportion by PB samples). (2) Followed by non-negative matrix factorization (NMF), or more precisely, Poisson Matrix Factorization (PMF), we can decompose the PB expression data (*Y*_pb_) into topic-specific gene expression (dictionary) matrix (*β*) and topic loadings/proportions for each PB sample (*θ*_pb_). The non-negativity constraint is natural to the gene expression count data and the additivity of factors generates biologically interpretable solutions. (3) Treating the dictionary matrix as a design matrix in a non-linear, non-negative regression problem, we can quickly deconvolve cell-level topic proportions (Fig. 1D). The resulting topic proportion matrix often results in markedly improved UMAP results (Fig. 1B vs. Fig. 1D). Moreover, the dictionary matrix *β* and two topic proportion matrices *θ*, estimated by PMF algorithm (Gopalan et al. 2014, 2015; Levitin et al. 2019), clearly exhibit modular structures in both sides. See Methods for technical details.

**Figure 1.**
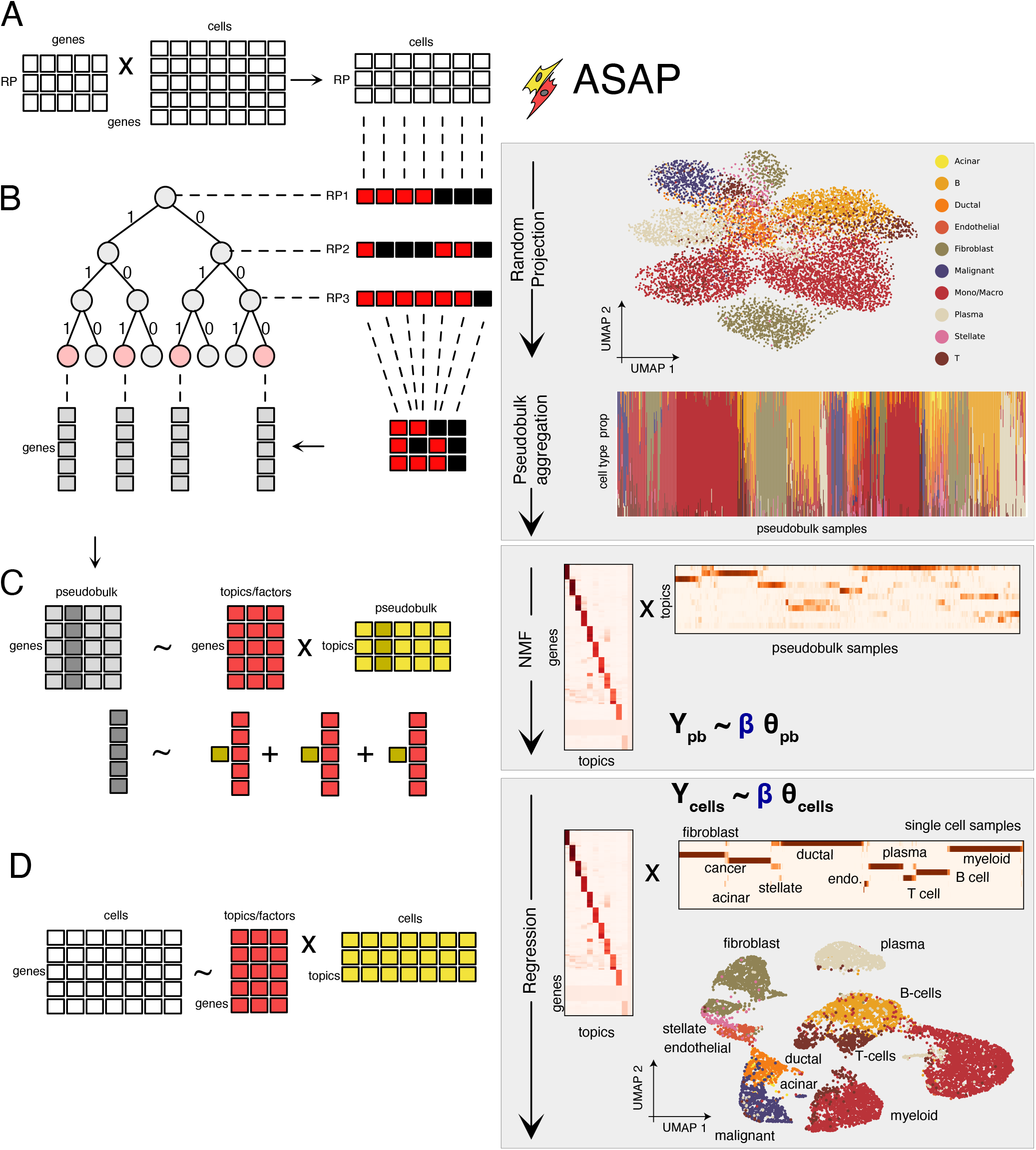
Schematic overview of ASAP framework. ASAP proceeds in three steps: collapsing cell count data into a manageable pseudo-bulk matrix (A), factorizing the pseudo-bulk matrix by Poisson matrix factorization (C), and regressing cell-level data onto topic space (D). For illustration purpose, we show intermediate results in processing pancreatic ductal adenocarcinoma data, consisting of 9,887 cells (Zhao et al. 2021a). **(A)** A raw gene expression matrix can be projected on the *r*-dimensional space of RP variables (right) by multiplying the observed cell count matrix (middle) with some univariate Gaussian matrix (left). **(B)** According to the sign of each random projection (RP) direction, switching to the left and right at each level, cells are sorted via a perfect binary tree with the depth= *r* (RP variables) and aggregated into pseudo-bulk samples. **(C)** A non-negative matrix factorization method decompose the pseudo-bulk data matrix into dictionary (gene by topic/factor) and factor loading matrix (sample by topic/factor). We represent each pseudo-bulk sample as a non-negative linear combination of non-negative topic-specific gene expression topics. **(D)** Treating the dictionary matrix as a design matrix, ASAP revisits the original gene expression vectors of many cells and regresses them on topic space with the same non-negative constraints on the regression coefficients, which can be used to identify cell-level clustering patterns (UMAP on the right).

### ASAP accurately estimates cell clusters

In single-cell data analysis, it is implicitly assumed that we collect a sufficient number of cells within each cell type, and these cells show more similar expression patterns within the same cell type than between different cell types. Some intrinsic, biological similarity measures can group cells of identical cell types/states. Had dimensionality reduction or latent embedding methods well-preserved cells’ biological similarity, clustering results based on the latent factors/topics should have been closely matched with the groups based on ground truth cell types. Hence, we evaluate the performance of different embedding methods in terms of the correspondence between cell groups found by clustering each method’s latent factors/topics and ground truth cell types.

We generated realistic simulation data treating cell type-specific gene expression profiles in cell sorting experiments (Schmiedel et al. 2018). We used the sorted bulk sequencing data instead of other single-cell data sets that are computationally annotated by some clustering methods. To gain insights into variability induced by sequencing depths and sample size, we varied the number of cells from 1,000 to 10,000, also varying the proportion of cell type-specific gene expression and background variation (see Methods). Since we are interested in comparing the estimated set membership (based on clustering) with actual cell type membership, we used three performance metrics widely used to evaluate clustering tasks: adjusted Rand index (ARI) (Rand 1971), normalized mutual information (NMI) (Vinh et al. 2010), and purity score (Liu et al. 2020; Kriebel and Welch 2022).

### Random projection well-preserves underlying cell type identities

We first wanted to understand the general behaviour of random projection operation. A similar analysis was conducted in previous studies, such as (Wan et al. 2020). As we increase the purity (*ρ*) level (Fig. 2A), patterns emerge, and cells form groups (Fig. 2B). Since visual inspection via UMAP is not rigorous enough, we evaluated the accuracy of Leiden clustering results (Traag et al. 2018) based on different configurations of latent factors–derived from PCA and RP with a different number of components/factors (Fig. 2C). Although the performance of RP data generally lags behind that of PCA by significant statistical margins, noting that PCA is more computationally intensive, we found that RP operation can be used as a good-enough-approximate method for the subsequent steps. Conversely, it also shows that RP alone is insufficient to capture a detailed view of cellular diversity, but the results of RP need to be refined.

**Figure 2.**
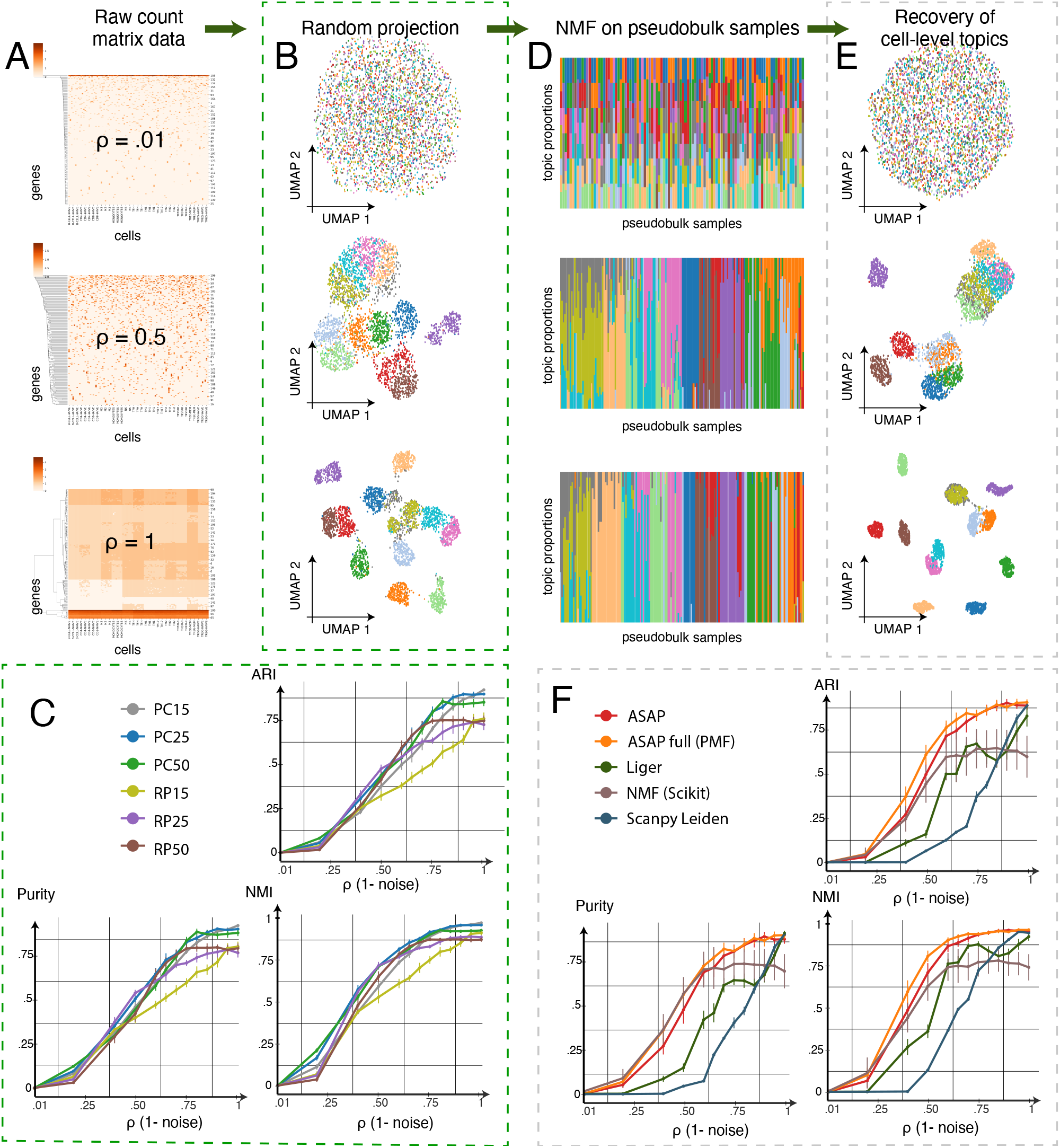
ASAP accurately estimates cell clusters. Extensive benchmark experiments confirm that our approximating method performs better or at least as well as existing methods. **(A)** We generated realistic simulation data treating cell type-specific gene expression profiles in cell sorting experiments (Schmiedel et al. 2018) as a gold standard while applying a different noise level, or conversely increasing proportion of cell-type-specific variance from *ρ*= .01 to *rho* = 1 (the top: *ρ* = 0.01; the middle: *ρ* = 0.5; the bottom: *ρ* = 1). **(B)** We can reduce dimension by multiplying the simulation count matrix with a random Gaussian matrix (see Fig. 1A and Methods). UMAP visualization shows how cell type-specific clusters emerge as we increase the proportion of cell type variance from the top (*ρ* = 0.01) to bottom (*ρ* = 1). Each dot represents a cell with its originating cell type (colour). **(C)** Cell-level topic regression refines cell type-specific clusters. We show the UMAP projections based on the final topic regression results. **(D)** We tested the impact of different choices of RP dimensions on cell type clustering performance. We compared clustering results derived from different embedding methods with the gold standard cell type annotations (cell-sorted data). We report three different metrics on the y-axes–ARI (adjusted Rand index, the left), NMI (normalized mutual information, the middle), and purity (the right) while varying the noise levels. The higher, the better. RP-*k*: Leiden clustering of cell-cell 15-nearest neighbour graphs built on *k*-dimensional random projection results. PC-*k*: Leiden clustering of cell-cell 15-nearest neighbour graphs built on top *k* principal components. **(E)** We compared the accuracy of ASAP’s final topic embedding results against other NMF-based methods, such as ASAP-full (Poisson factorization directly applied to full data), Liger (Welch et al. 2019; Kriebel and Welch 2022), NMF function, `sklearn.decomposition` .NMF, implemented in SciKit-Learn library (Pedregosa et al. 2011), and a standard clustering pipeline implemented in `scanpy` library (Wolf et al. 2018). X-axis: the proportion of cell type variance; y-axes: ARI (adjusted Rand index, the left), NMI (normalized mutual information, the middle). The higher, the better.

### Poisson Matrix Factorization methods achieve a higher level of robustness

As a refinement step, we used PMF (Gopalan et al. 2014, 2015; Levitin et al. 2019)–applied to PB data matrix (`ASAP`) or full data matrix (`ASAP-full`/PMF). So, it was natural to compare with other NMF-based methods (Fig. 2E-F), which include Liger (Welch et al. 2019; Kriebel and Welch 2022), a standard NMF method (`sklearn.decomposition.NMF`) in scikit-learn library (Pedregosa et al. 2011), along with a conventional Leiden clustering pipeline implemented in `scanpy` library (Wolf et al. 2018). All the methods were summarized into the clustering results, and cell memberships were tested against the gold standard (see Methods for details). ASAP, Liger, and Scanpy’s clustering method naturally incorporates Leiden clustering as a final step. For the Leiden-based methods, we varied the resolution parameters (0.1, 0.25, 0.5, 0.75, and 1.0); our results were, albeit, invariant to the fine-tuning of Leiden algorithm. For the `scikit-learn` NMF, we simply used the standard k-means algorithm implemented in the same library.

We found that the PB-based ASAP and ASAP-full clearly outperformed other NMF and clustering methods with statistically significant margins, especially in settings with moderate and high noise levels (Fig. 2F), suggesting that the PMF algorithm, which both ASAP and ASAP-full were built on, is more robust to other NMF algorithms. PMF is a Bayesian hierarchical model, resolves its parameters by posterior inference and often yields more robust results than optimization-based algorithms. Perhaps a more important aspect of this benchmark result is that our ASAP method based on PB data is as competent as PMF fitted on full data (the red and Orange cures in Fig. 2F).

### ASAP is a scalable method with little memory footprint

Along with the performance metrics (Fig. 2), we also measured the run time (in minutes) and memory footprint (in gigabytes/GB) of each method in the same workstation. We allocated the same resources for all the experiments, fixing the available memory to 32GB and eight CPU cores. As long as the allocated memory permits, we varied the number of cells from 1,000 to 100,000. In the first batch of experiments, varying the number of cells from 1,000 to 10,000, ASAP-full (vanilla PMF) often took the most time (Fig. 3A) and emended high memory to keep track of multiple copies of variational parameters (Fig. 3B), followed by `Liger` and `scikit-learn` NMF methods. However, ASAP was consistently fast enough to get the results within a minute and kept the peak memory usage under 2GB until the 10k cell experiments.

**Figure 3.**
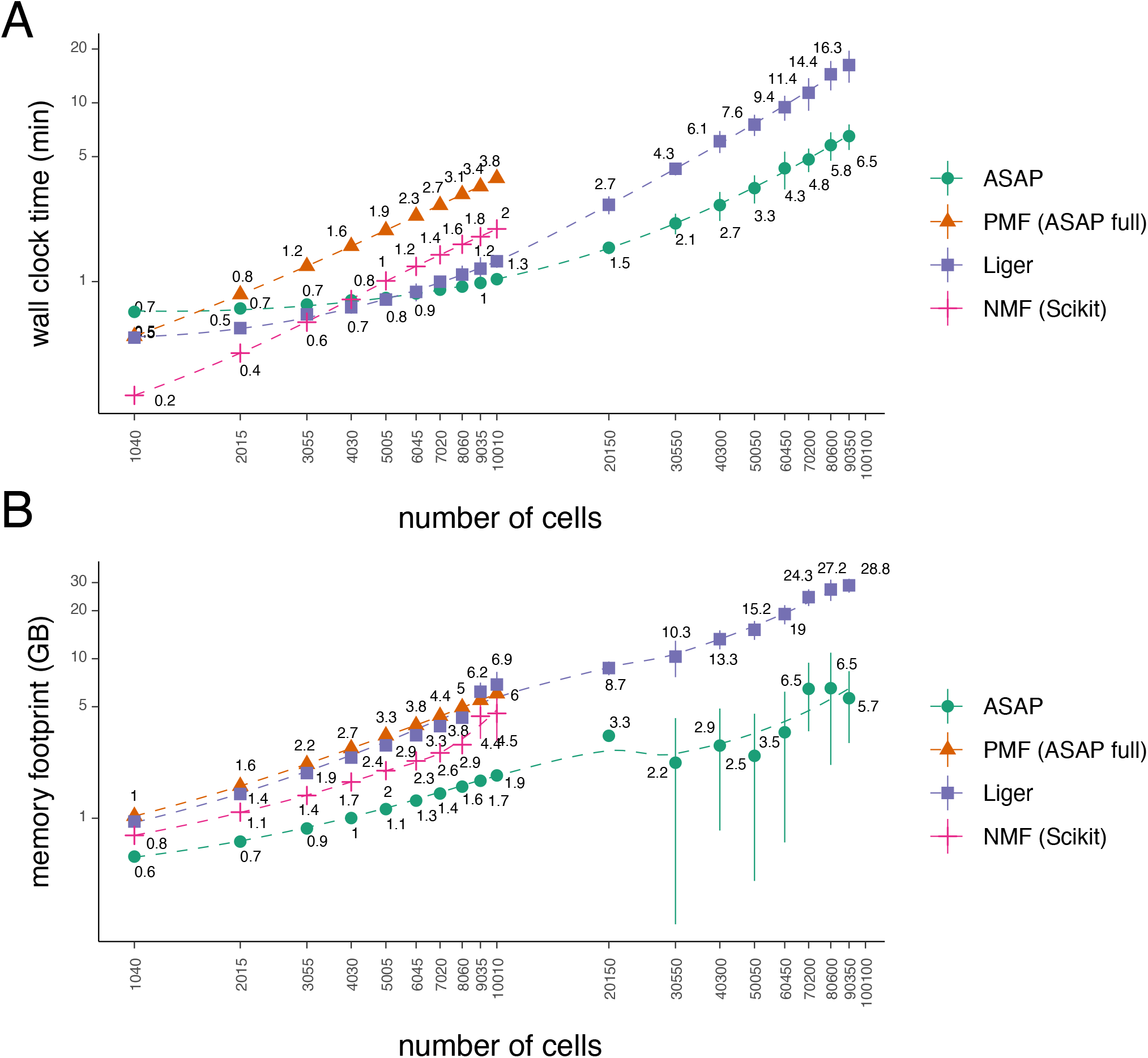
ASAP achieves scalability in terms of computation time and memory footprint. The plot shows **(A)** the elapsed time (in minutes) and **(B)** peak memory usage (in GB) by NMF-based methods with different numbers of cells ranging from 1000 to 100,000 as a function of the number of cells.

In the second batch, we focused on Liger and ASAP models, varying the number of cells from 20,000 to 100,000. Here, ASAP showed significantly faster runtime and lower memory usage. Liger already peaked its memory usage near 32GB with 100,000 cells. Therefore, it was not included in further experiments. We also conducted further experiments with ASAP alone to measure runtime and memory usage and found them to scale linearly with the number of cells (Supplementary figure).

### ASAP recapitulates known cell-type-specific gene programs in real-word data sets

We applied ASAP to three different large-scale scRNA-seq data sets to demonstrate that our method is scalable and accurate enough to recapitulate well-established cell-type-specific gene expression patterns. We considered the following date sets as representative examples: (1) peripheral blood mononuclear cells (PBMC) (Hao et al. 2021) profiled on 161,764 cells and 20,729 genes (Fig. 4A-C);(2) breast cancer mammary tissues (Wu et al. 2021) of 20,265 genes expressed on 100,064 cells (Fig. 4D-E); (3) the first draft of Tabular *Sapiens* project (Tabula Sapiens Consortium et al. 2022) (Fig. 4F-G), consisting of 483,152 cells and 58,604 genes/transcripts.

**Figure 4.**
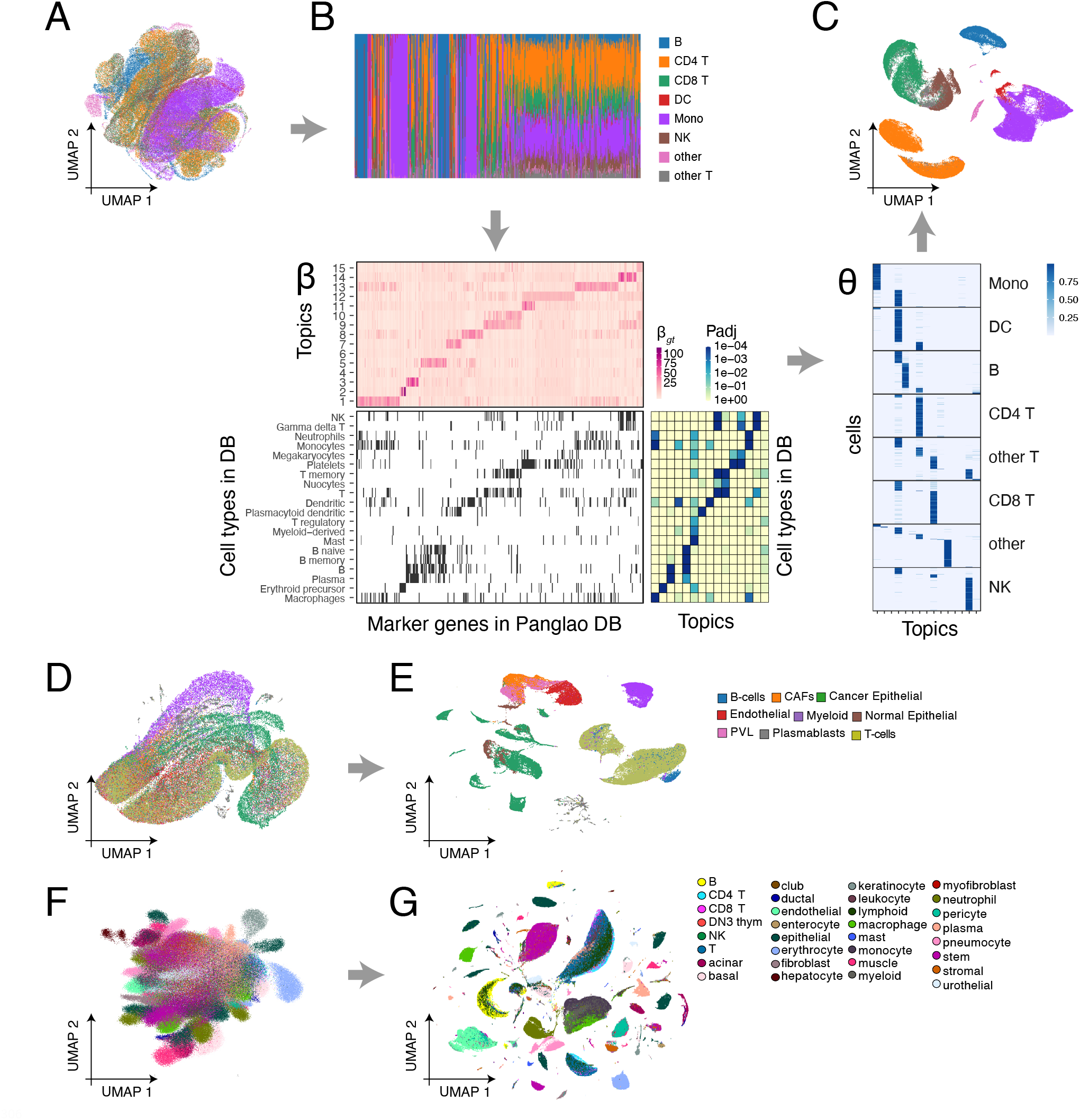
ASAP accurately annotates cell types in real-world large-scale data. *Date sets:* peripheral blood mononuclear cells (PBMC) data (Hao et al. 2021) (A-C), breast cancer mammary tissues (Wu et al. 2021) (D-E), Tabular *Sapiens* data (F-G) (Tabula Sapiens Consortium et al. 2022). We show the three steps of ASAP applied to the PBMC data (A-C). **(A)** Step 1: UMAP drawn directly from the RP results. The colours represent different cell types previously annotated (level 1) by the original authors (Hao et al. 2021). ^*^**(B)** Step 2: Pseudobulk data (top) along with the PMF result–a dictionary matrix *β* (bottom). We compared the gene activities in the *β* matrix with the known cell type marker genes (Franzén et al. 2019). **(C)** Step 3: Fast cell-level regression analysis to recover topic proportions of all the cells. UMAP (top) drawn from the topic proportion results *θ* (bottom). **(D)** UMAP drawn directly from the RP results of the breast cancer data (Wu et al. 2021). **(E)** UMAP based on the final topic proportion results of ASAP. The colours represent different cell types based on the original study. **(F)** UMAP directly drawn from the RP results of the Tabular Sapiens data (Tabula Sapiens Consortium et al. 2022). **(G)** UMAP based on the final topic proportion results of ASAP. The colours represent different cell types based on the original study (the broad level).

ASAP’s dictionary matrix *β* (topic-specific gene expressions) strongly enriches the known cell type marker genes,`PanglaoDB` (Franzén et al. 2019), within the same topics (bottom, Fig. 4B); hence, the resulting cell topic proportions clearly differentiate cells based on the actual cell type annotations (Fig. 4C). We performed gene set enrichment analysis (GSEA) within each topic by `fgsea` (Korotkevich et al. 2021), which provides faster and more accurate approximation of traditional GSEA method(Subramanian et al. 2005). Interestingly, the same level of resolutions was not observed in the first RP step (Fig. 4A). Many PB samples were rather heterogeneous, mixing multiple cell types (top, Fig. 4B).

Considering that no label information was given to the factorization step, our intermediate results demonstrate that the non-negative matrix factorization step is indeed critically important, and the previous PB construction captures necessary information while compressing data complexity drastically. In the other real-world data analysis, we also observed that PB construction provided rough ideas about cell-type-specific patterns (Fig. 4D, F); the subsequent PMF steps refine topic-specific gene activities to result in high-resolution cell-type annotations in the end (Fig. 4E, G). We also noted that the topics do not always agree with the cell types, which are often best characterized by cell type-specific surface markers and transcription factors. For instance, the topics of the breast cancer data may implicate finer resolution of cancer subtypes (Fig. 4E), or the different cell types could be strongly affected by shared topic-specific genes (Supplementary Fig. 1). For a large study, such as Tabular Sapiens, cell type annotations are often manually curated by different researchers with different domain knowledge, e.g., dividing the jobs tissues-by-tissue. Nonetheless, we found the original annotations generally correspond to topics (Supplementary Fig. 2) and are well-separated in the topic space (Fig. 4G).

### Pseudobulk samples correspond to bulk expression samples

We had expected that PB samples would behave similarly to cell-type-sorted bulk samples. However, the previous real-world PB data showed quite a level of heterogeneity, mixing multiple cell types– especially those closely related cell types, such as CD4+ and CD8+ T-cells. Such results also show that RP alone is insufficient to investigate fine-resolution cellular diversities.

To better understand the characteristics of PB samples, we investigated single-nucleus RNA-seq (snRNA-seq) (Eraslan et al. 2022) and bulk RNA-seq data (GTEx Consortium 2020) profiled by Genotype-Tissue Expression (GTEx) consortium. This snRNA-seq data set provides gene expression vectors on 209,126 cells sampled from 16 individuals across ten tissue types, for which the GTEx consortium also provides publicly-accessible data (GTEx Consortium 2020).

We first analyzed the snRNA-seq matrix with ASAP and obtained its dictionary *β* matrix (gene by factor). As can seen in the GSEA against `PanglaoDB` (Franzén et al. 2019) marker gen sets by the `fgsea` method (Korotkevich et al. 2021; Subramanian et al. 2005), these topics are significantly associated with known cell types (bottom, Fig 5A). We also found that our unsupervised learning can recover the broad cell-type annotations called by the original study (Fig. 5B). Interestingly, most cell types are shared across multiple tissues, yet there still exist several tissue-specific cell types, such as multiple epithelial cell types and myocytes, perhaps, more specialized in tissue-specific environments.

**Figure 5.**
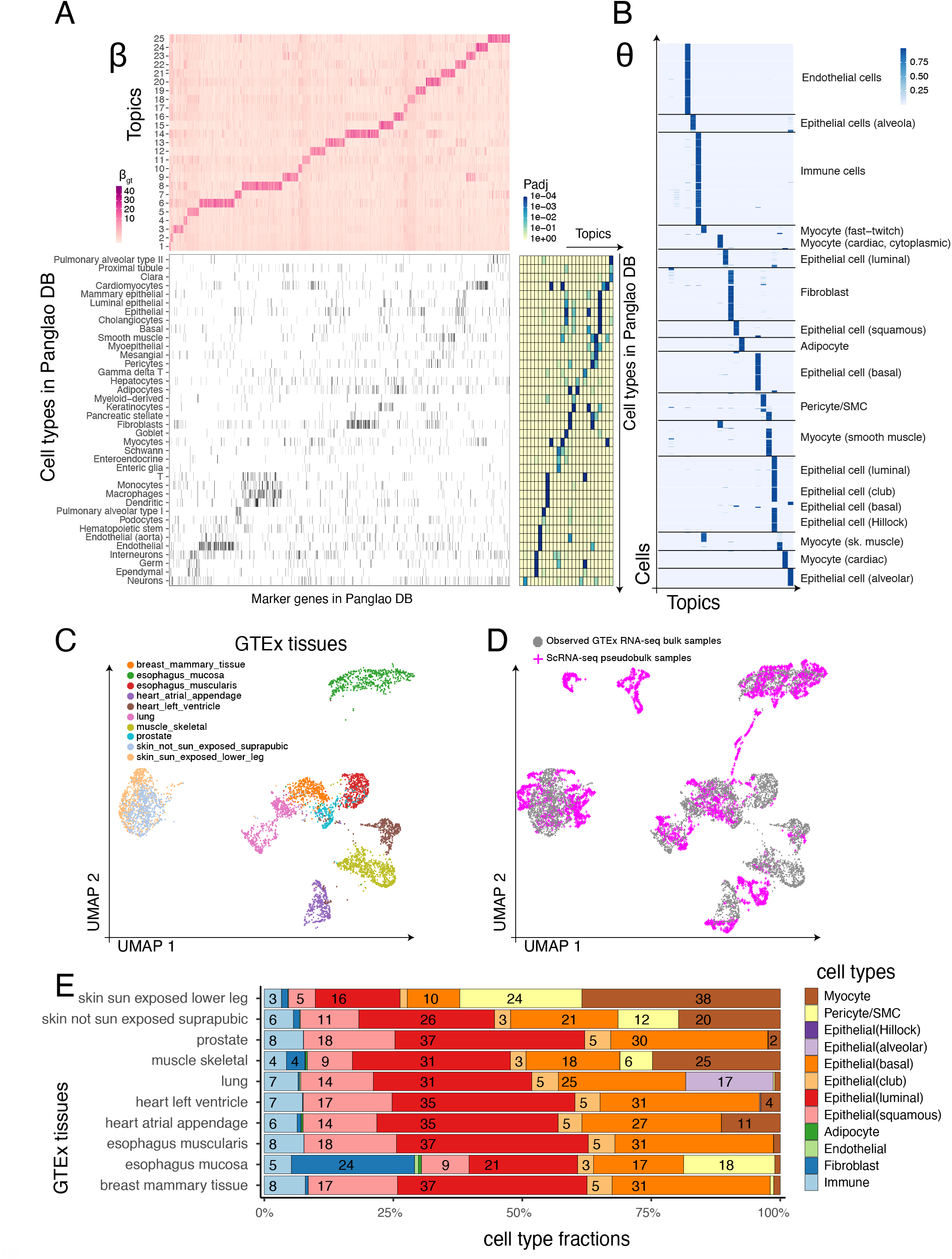
ASAP deconvolves cell type-specific gene programs in tissue-level data in joint analysis with snRNA-seq data. *Date sets:* Genotype Tissue Expression project’s single-nucleus RNA-seq data (Eraslan et al. 2022), GTEx project’s bulk RNA-seq data (GTEx Consortium 2020) (D-E). **(A)** Topic-specific gene activity matrix *β* trained on the pseudobulk data constructed in the GTEx snRNA-seq data (Eraslan et al. 2022) shows strong enrichment of known cell types in PanglaoDB (Franzén et al. 2019). *β*: topic (rows) by gene (columns). (B)The patterns of ASAP’s topic proportion estimates agree *θ* with the original annotations. *θ*: cell (rows) by topic (columns). **(C)** UMAP embedding of GTEx bulk data, only including tissue types present in the snRNA-seq data. Before the UMAP, we projected the bulk data onto the snRNA-seq topic space. The colours represent different tissues of origin. **(D)** The snRNA-seq pseudobulk samples (magenta) are overlaid on the same snRNA-seq topic space where the bulk samples (gray) are projected previously. **(E)** Cell type proportions estimated by the joint analysis of GTEx snRNA-seq and bulk data. X-axis: cell type fractions; y-axis: different tissue types; the numbers in the segments: fraction of cell types in percentage.

Using the dictionary matrix *β*, we projected the bulk RNA-seq data onto the snRNA-seq topic space (Fig. 5C-D); then, we overlaid the snRNA-seq PB data on top of the same coordinates. GTEx tissues are generally well-separated from one another (Fig. 5C), suggesting different cell type factions as a primary axis of variation. Moreover, we found that the PB samples overlap with these bulk samples to a large degree. Such good coverage compelled the follow-up experiment to assess whether the projected coordinates could be a basis for cell type deconvolution step. Simply, we were able to query one hundred neighbouring snRNA-seq cells for each bulk sample and count the frequency of each cell type within the neighbours.

The neighbouring cell type fractions can be interpreted as approximate deconvolution profiles (Fig. 5E). Unlike marker gene-based deconvolution methods, no prior knowledge was elicited, and reference cell type profiles were adaptively selected for each bulk sample. Our results suggest several expected patterns. Substantially higher fractions of myocytes were found in the sun-exposed side of skin and skeletal muscles (brown colours). A special type of alveolar epithelial cells was specifically found in the lung tissues. Immune cells are present in all the tissues but much less than other cell types. The two types of esophagus tissues, histologically distinctive, are markedly different in terms of the cell type fractions. We found a more diverse mixture in the esophagus mucosa, containing a much higher fraction of fibroblasts and pericytes.

## Discussion

We propose a novel approximation method that can quickly identify topic structures embedded in large-scale single-cell data. Our method builds on the rationale that the distance between two high-dimensional data points can be preserved in low-dimensional space constructed by random with a high probability (Johnson and Lindenstrauss 1984; Bingham and Mannila 2001; **???**; Dasgupta and Freund 2008)f. Based on this foundation, a similar strategy has been used to speed up a clustering procedure (Wan et al. 2020); however, the quality of RP-based clustering analysis demands multiple random projections and clustering steps, and the resolution of clustering results are limited and unable to delineate a subtle difference between similar cell types; hence, the benefit of reduced computation time and peak memory can be diminished, and clustering results are often left less interpretable and require further *post hoc* analysis. Although our approach begins with a similar random projection, our key idea focuses on building a randomly sorted pseudobulk data matrix. In realistic benchmark analysis based on sorted bulk RNA-seq profiles, we demonstrate that learning a topic model on randomly sorted pseudobulk outperforms unsupervised cell-type annotation tasks. Unlike clustering results, our topic model parameters are immediately interpretable and directly comparable against known marker gene databases. Furthermore, we show that our approach resembles a data-generating scheme of bulk RNA-seq data, and the inference results are easily transferable for deconvolution analysis.

Our approach can be extended in many different ways. We can intuitively apply RP operations to reduce feature space for epigenomic, genomic, and other types of sparse matrix data. Although aggregating count data within a PB sample generally works robustly, we can combine batch effect adjustment and advanced depth normalization methods to further improve the quality of the resulting PB data. If needed, multiomics data integration can take place at a PB level followed by nearest neighbour matching (Haghverdi et al. 2018; Hie et al. 2019a; Polański et al. 2019) or even optimal transport (Demetci et al. 2022). Technically, any machine learning algorithm can be used to learn a topic-specific feature frequency matrix. A deep learning-based generative model can be more efficiently trained by alternatively applying RP and PB steps; proper modelling that incorporates prior knowledge of cell-type-specific marker genes and dependency structures will also substantially improve inference results (Elyanow et al. 2020; Townes and Engelhardt 2022).

## Methods

### Probabilistic topic modelling by ASAP

#### Goal: topic modelling for single-cell data analysis

Single-cell transcriptomic profiling results in a count matrix 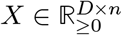of *n* cells where each column vector lies in *D*-dimensional space. The dimensionality *D* corresponds to the number of features, such as genes and non-coding RNA molecules. Each element *X*_*i*_ (*j* ∈ [*n*]) quantifies how many reads were mapped onto a gene *i* within a cell *j*; hence, each value is non-negative by nature and generated by a counting process. Throughout this work, we will indicate a gene by *i* ∈ [*D*], denote a cell/sample by *j*, and use *k* ∈ [*K*] for a topic index. In total, we have *D* features and *n* cells,for each cell. and the goal is to estimate *K* topics, (*β*_1_, …, *β*_*K*_), and the corresponding topic proportion vector,(*θ*_1_, …, *θ*_*n*_),for each cell.

Given that each cell *j*, the column vector **x**_*j*_, sits on a topic space, *k* ∈ [*K*], we can assign a topic proportion of a topic *k* to the cell *j*, namely *θ*_*jk*_ values, to quantify which and how much each topic affected each cell. Defining a matrix *β* with each element *β*_*ik*_ for an expected expression value for a gene *g* on a topic *k*, we assume that an observed gene expression matrix *X* was generated by a linear combination of the two matrices–*β* and *θ*. More precisely, we have

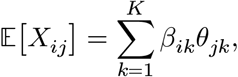

where the equality holds proportionally up to some scaling factor. The goal is to estimate the unknown gene topic (program) dictionary (*β*) and the topic proportion matrix (*θ*) provided with an observed gene expression matrix *X* and the fact that distinctive gene expression programs/topics exist and are expressed differently according to each cell’s topic representation.

#### Poisson matrix factorization

Poisson matrix factorization (PMF) model (Gopalan et al. 2014, 2015; Levitin et al. 2019) treats the same non-negative factorization problem as Bayesian inference problem, assuming that each element of the data matrix, *X*_*ij*_, was generated by the following generative model:

1. Sample topic *k*-specific gene/feature expression: *β*_*ik*_ ∼ Gamma (*a*_0_, *b*_0_).
2. Sample each cell/sample’s activity/loading on a topic *k*: *θ*_*jk*_ ∼ Gamma (*a*_0_, *b*_0_)
3. Constitute the sampling rate *λ*_*ij*_ for gene/feature *i* in each cell/sample *j* as a linear combination of the two parameters, namely *λ*_*ij*_ = ∑_*k*_ *β*_*ik*_ *θ*_*jk*_.
4. Sample gene expression for a gene *i* in a sample *j* from Poisson distribution: *X*_*ij*_ ∼Poisson(*X*_*ij*_ |*λ*_*ij*_)

More precisely, we define the probability density function (PDF) of Gamma(*λ*|*a, b*) as

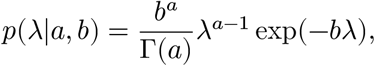

and the PDF of Poisson(*X*|*λ*) as

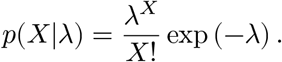

*Remark*: The scale of the *β* and *θ* parameters are arbitrary, and they can adapt to each other, e.g.,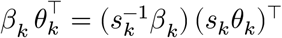. The lack of identifiability in scaling factors brings about interesting equivalence relationship between PMF and conventional multinomial topic model likelihood (Carbonetto et al. 2021). It is straightforward to translate the PMF parameters into the conventional topic model parameters by adjusting the scales of *β* and *θ* parameters adaptively, such that ∑_*k*_ *β*_*jk*_ = 1 and ∑_*k*_ *θ*_*jk*_ = 1.

### Detailed steps of ASAP algorithm

#### Step 1. Random projection (RP) and pseudo-bulk (PB) construction

A full *D* × *n* data matrix *X* can be sparse and too big to be read and densified in local memory. In the first step,each column/cell vector **x**_*j*_ is compressed to a smaller, manageable *d* × *n* matrix *Q* by randomly projecting the total data matrix *d* times onto *d*-dimensional space (*d* ≪ *D*). For each random projection (*k* = 1, …, *d*), we use isotropic *D*-dimensional Gaussian vector **r**_*k*_ and take the dot product with each cell vector,namely,

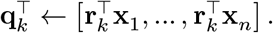

To ensure that each RP vector is independent of the others, we perform (economical) singular value decomposition (SVD) of the *Q* matrix and construct orthogonal *d* × *n* RP matrix 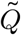. After applying row-wise standardization, for each cell *j* ∈ [*n*], we can quickly convert 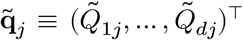 to a vector of *d* binary codes by setting each *k*-th coordinate to 1 if the 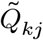 is positive; otherwise, it is 0 (Fig. 1B). This binary code vector can be used as a binary sorting tree, literally converting a binary number to a decimal one, with which we can quickly scatter cells into 2^*d*^ buckets (leaf nodes). In each bucket/leaf node, we can incrementally aggregate expression vectors to a PB sample, and each PB vector **y**_*l*_ is later normalized to have uniform sequencing depths (*l* = 1, …, 2^*d*^). For simplicity,we ignore empty buckets; hence, the resulting PB matrix can have at most 2^*d*^columns/samples.

- Construct a raw RP matrix *Q*_*d*×*n*_ ← *R*_*d*×*D*_*X*_*D*×*n*_.
- Orthogonalize the *Q* matrix and build 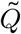 by SVD and standardization
- Make a binary *B* matrix by setting each element

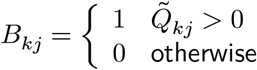

where *k* = 1, …, *d* and *j* = 1, …, *n*.
- Convert this *B* matrix to a leaf node (bottom bucket) membership matrix:

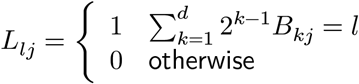

where *k* = 1, …, *d* and *j* = 1, …, *n*.
- Aggregate cells within each leaf node into a PB sample:

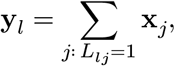
- Normalize **y**_*l*_ ← **y**_*l*_/‖**y**‖_1_ × 10^4^.

#### Step 2. Variational inference of Poisson matrix factorization model

Next, we decompose the *d* × *L* PB matrix *Y* by PMF model (*L* ≤ 2^*d*^), modelling each PB data point as Poisson distribution with the rate parameter that can be decomposed into dictionary and loading matrices (see the previous subsection for details). However, the log-likelihood,

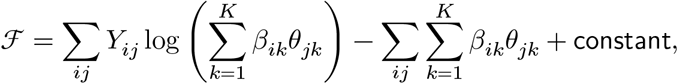

is non-conjugate with the Gamma distributions of the *β* and *θ* parameters. Following the previous variational inference algorithm (Gopalan et al. 2014, 2015; Levitin et al. 2019), we need to introduce auxiliary variables to construct evidence lower bound (ELBO) under the mean-field variational approximation (using Jansen’s inequality):

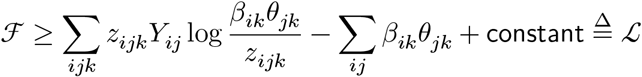

This ELBO now permits the following closed-form update equations for the variational distributions.

- Inference of the local/sample-specific topic proportions:

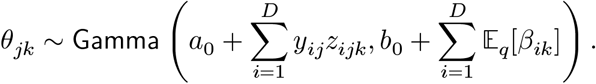
- Inference of the global/dictionary parameters:

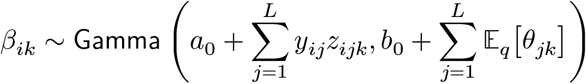
- Inference of the auxillary variable:

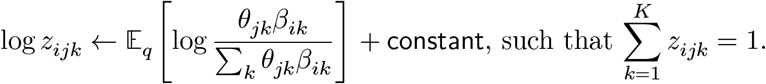

#### Step 3. Prediction using latent variable factorization

Finally, given that we have the expected dictionary matrix, namely 𝔼[*β*] and 𝔼[log *β*], we revisit full single-cell data and recover cell-level topic proportion vectors, namely *θ*_*j*_ = (*θ*_1*j*_, …, *θ*_*kj*_, …, *θ*_*Kj*_)^⊤^ for each cell *j* = 1, …, *n*. We could use the same variational inference algorithm, only skipping the update for *β* the parameters, but it would involve full-scale inference of the auxiliary variables and not be scalable in practice. Instead, we form a different approximation of the ELBO and calibrate the *θ* parameters by solving a massive array of regression problems:

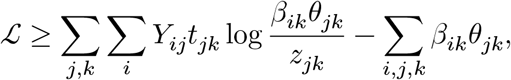

where ∑_*k*_ *t*_*jk*_ = 1 for all *j*. Letting *ρ*_*jk*_≜ 𝔼[*z*_*jk*_] under the same variational approximation, we can derive two closed-form update equations. Firstly, we have

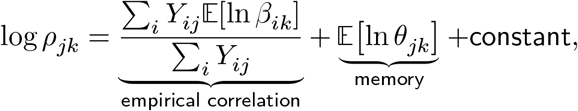

where ∑_*k*_ *ρ*_*jk*_ = 1, and have the variational

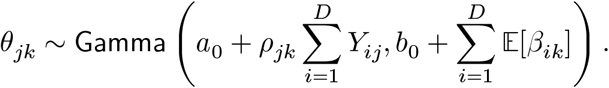

## Simulation

### Simulation scheme

We generated a single-cell dataset from cell-type-sorted bulk RNA-seq reference data from the DICE (Database of Immune Cell Expression, Expression quantitative trait loci and Epigenomics) project (Schmiedel et al. 2018) to evaluate the performance of ASAP and other methods with gold standard annotations. We considered the population of thirteen cell types, including CD4+ T cells, CD8+ T cells, NK cells, B cells, and monocytes. We use a well-established data-generating framework implemented in the `scDesign2` package–a simulation method based on a Gaussian copula-based sampling scheme (Sun et al. 2021). Gene-gene correlation structures are accurately captured in data simulated by a copula method, and the non-parametric nature of its density estimation step is more suitable for simulation experiments with little modelling assumption. Mathematical details can be found in the original paper (Sun et al. 2021), but briefly, we simulated our data as follows.

For each cell type *t*, we generate a single-cell data matrix *Y*^(*t*)^based on a continuous version of the transformed data as follows:

1. Estimate gene-level mean *μ* and covariance between genes Σ for each cell type *t*.
2. For each cell *j*, we generate bootstrapped copula **y**_*j*_:
  - Sample a Gaussian vector **z**_*j*_ ∼ 𝒩(*μ*_*t*_, Σ_*t*_).
  - Construct a stochastic version of the reference gene expression vector **s**_*j*_ by sampling with replacement.
  - Sort the bootstrapped gene expressions *S*_*gj*_, i.e., *S*_(1)*j*_ < *S*_(2)*j*_ < ⋯ < *S*_(*D*)*j*_.
  - Assign each gene *g*’s gene expression *Y*_*gj*_ in two steps:
    – Identify the ascending order *r* for *Z*_*gj*_
    – Take the bootstrapped *S*_(*r*)*j*_ and set *Y*_*gj*_ to this value.
3. Repeat the above steps until we sample the desired number of cells.

Next, we simulated a separate single-cell data matrix from a null model that captured the global gene expression pattern across all cell types in the bulk samples. To account for the background and cell-type-invariant patterns, we generate the null data 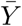 by sampling a gene vector 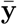 from a multinomial distribution with the empirical gene frequencies, ignoring cell-type-identities. Depending on the *ρ* ∈ (0,1) value, we differently mix the cell-type-specific foreground and the background signals: 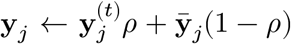. Both foreground and background data were normalized to have the same sequencing depth.

## Notes

### Competing Interest Statement

The authors have declared no competing interest.

